# The transcriptomic profiling of COVID-19 compared to SARS, MERS, Ebola, and H1N1

**DOI:** 10.1101/2020.05.06.080960

**Authors:** Alsamman M. Alsamman, Hatem Zayed

**Affiliations:** Department of Genome Mapping, Molecular Genetics and Genome Mapping Laboratory, Agricultural Genetic Engineering Research Institute, 9 Gamaa St. 12619, Giza, Egypt; Department of Biomedical Sciences College of Health Sciences, QU Health, Qatar University, Doha, Qatar

**Keywords:** COVID-19, Ebola, transcriptome, protein-protein interaction, gene expression, pandemics

## Abstract

COVID-19 pandemic is a global crisis that threatens our way of life. As of April 29, 2020, COVID-19 has claimed more than 200,000 lives, with a global mortality rate of ~7% and recovery rate of ~30%. Understanding the interaction of cellular targets to the SARS-CoV2 infection is crucial for therapeutic development. Therefore, the aim of this study was to perform a comparative analysis of transcriptomic signatures of infection of COVID-19 compared to different respiratory viruses (Ebola, H1N1, MERS-CoV, and SARS-CoV), to determine unique anti-COVID1-19 gene signature. We identified for the first time molecular pathways for Heparin-binding, RAGE, miRNA, and PLA2 inhibitors, to be associated with SARS-CoV2 infection. The *NRCAM* and *SAA2* that are involved in severe inflammatory response, and *FGF1* and *FOXO1* genes, which are associated with immune regulation, were found to be associated with a cellular gene response to COVID-19 infection. Moreover, several cytokines, most significantly the *IL-8*, *IL-6*, demonstrated key associations with COVID-19 infection. Interestingly, the only response gene that was shared between the five viral infections was *SERPINB1*. The PPI study sheds light on genes with high interaction activity that COVID-19 shares with other viral infections. The findings showed that the genetic pathways associated with Rheumatoid arthritis, AGE-RAGE signaling system, Malaria, Hepatitis B, and Influenza A were of high significance. We found that the virogenomic transcriptome of infection, gene modulation of host antiviral responses, and GO terms of both COVID-19 and Ebola are more similar compared to SARS, H1N1, and MERS. This work compares the virogenomic signatures of highly pathogenic viruses and provides valid targets for potential therapy against COVID-19.

## Introduction

COVID-19 is caused by severe acute respiratory syndrome coronavirus 2 (SARS-CoV-2). As of April 25, 2020, the COVID-19 pandemic has spread to more than 200 countries and territories with about 3 million confirmed cases and ~ 7 % mortality (WHO 2020).

SARS-CoV-2 belongs to the *Coronaviridae*, of this family the severe acute respiratory syndrome coronavirus (SARS-CoV/SARS) and the Middle East Respiratory Syndrome Coronavirus (MERS-CoV/MERS). In 2002 and 2012, SARS and MERS were associated with ~8,000 cases and ~2,500 cases with a case fatality rate of ~10% and ~36%, respectively. As with previous coronaviruses, there are no specific antivirals or approved vaccines available to control SARS-CoV-2, SARS, or MERS, where only conventional control measures, including travel restrictions and patient isolation, could stop or slow down their social impact (Hoffmann et al. 2020). The full-length genome of SARS-CoV-2 revealed 87.99% sequence similarity with the bat SARS-like coronavirus and 80% identity nucleotide with the original SARS epidemic virus (Tan et al. 2020).

The current outbreak of SARS-CoV-2 virus is very similar to Ebola virus disease (EVD) and influenza A virus subtype H1N1 outbreaks in 2009 and 2013–2016. Similar to SARS-CoV-2, the main reservoir for EVD is considered to be bats where the magnitude of its outbreak was unprecedented, with > 28 500 reported cases and > 11 000 deaths in West Africa (Vetter et al. 2016). On the other hand, swine-origin influenza (H1N1) spread rapidly throughout the world, leading the world health organization (WHO) to declare a pandemic on June 11, 2009 (Girard et al. 2010). The virus was determined to be an H1N1 virus clinically and antigenically identical to seasonal influenza viruses in humans and closely similar to swine circulating viruses (Schnitzler and Schnitzler 2009).

Fatigue, fever, dry cough, myalgia, and dyspnoea are the most common symptoms at the onset of SARS-CoV-2 infection and the less common symptoms were nausea, headache and gastrointestinal symptoms (Song et al. 2020). The most common indicator of radiological detection was the bilateral ground-glass or patchy opacity. Some of the patients had lymphopenia and eosinopenia. Blood eosinophil levels positively correlates with the lymphocyte levels following hospital admission in severe and non-severe patients. Severe patients were associated with substantially higher levels of D-dimer, C-reactive protein, and procalcitonin relative to nonsevere patients (Zhang et al. 2020). A typical biological response for the different viral infections may be identified, whereas some particular genes are dysregulated during the infection with specific viruses. Such response may have a major impact on the ability of the host to mount an adaptive host response. For instance, both MERS and SARS-CoV induced a similar activation of pattern recognition receptors and the interleukin 17 (IL-17) pathway (Josset et al. 2013).

Only victims of crime may describe their perpetrators, only they can use the best words to convey their experience. In the case of coronavirus, our own transcriptome is the victim, so we need to listen to its comprehensive description. We conducted an extensive study of the transcriptomic response of SARS-CoV-2. We located common and specific differential expressed genes to SARS-CoV-2 that are shared with SARS-CoV, MERS-CoV, H1N1, and Ebola. We performed chromosomal location, gene ontology and protein-protein interaction for such genes in order to understand SARS-CoV-2 unusual high infection rate and mortality. These profiles could be used to better understand the relationship between virus and host and to detect distinct responses to the expression of the host gene. This could have an impact on *in vivo* pathogenesis and could guide therapeutic strategies against the evolving virus.

## Material and Methods

### Datasets

The gene expression data of COVID-19, Ebola, H1N1, MERS-CoV, SARS-CoV have been retrieved from NCBI-GEO archive (Barrett et al. 2009), with ID GSE147507,GSE86539, GSE21802, GSE100504, and GSE17400, respectively. These data are based on Affymetrix human genome gene chip sets and Illumina NextSeq 500, revealing the gene expression profiles of *in vitro* and *in vivo* infections (**Table S1**).

### Data Analysis

The identification of the differentially expressed genes (DEGs) in the transcription profile was analyzed using GEO2R tool (Barrett et al. 2012) and differential expression analysis using DESeq2 and DEApp (Li and Andrade 2017) using default parameters. DEGs were characterized for each sample (p-value < 0.01) and were used as query to search for enriched biological processes. Gplot package in R was used to construct the gene expression heatmaps. The evaluation of the protein interactions and gene ontology (GO) enrichment was conducted with the STRING database (Szklarczyk et al. 2016). Cytoscape software has been used to visualize the structures of protein-protein networks (Shannon et al. 2003). The Circos software (Krzywinski et al. 2009) was used to represent gene expression and gene ontology analysis of the host response to viral infections based on human genome data (GRCh38).. The online tool Draw Venn Diagram (http://bioinformatics.psb.ugent.be/webtools/Venn/) was used to sketch a Venn diagram demonstrating some analysis information.

## Results

We investigated the unique transcriptomic gene expression signature that was induced by COVID-19 (GSE147507) compared to Ebola (GSE86539), H1N1 (GSE21802), MERS-CoV (GSE100504), and SARS-CoV (GSE17400), using the DEGs in transcriptomic profiles. The chromosome location of these DEGs sets are categorized according to the viral infection in **Figure 1**, in addition to the significant involvement of these genes in the response of different viral infections, based on pvalue (**Figure 1A-1F** and **Table S2)**. We identified 358 DEGs with a significant associated p-value < 0.01to COVID-19. Of these, *SAA2*, *CCL20*, *IL8* were highly significant (**Figure 1B** **and Table S2**). The analysis of gene enrichment of DEGs associated with the host response to COVID-19 highlighted several GO terms (**Figure 2**), including leukocyte activation, humoral immunity, myeloid cell activation, neutrophil activation, tuberculosis response, and miRNA involvement in the immune response. Additionally, GO terms that are correlated with cell death are highly consistent (**Figure 2C and 2E**). GO cytokine response terms, IL-17 signaling pathway, NF-kB ssignaling, TNF signaling pathway, and NF-kappa B signaling are among the most significant pathways associated with COVID-19 (**Figure 2B**).

**Figure 1:**
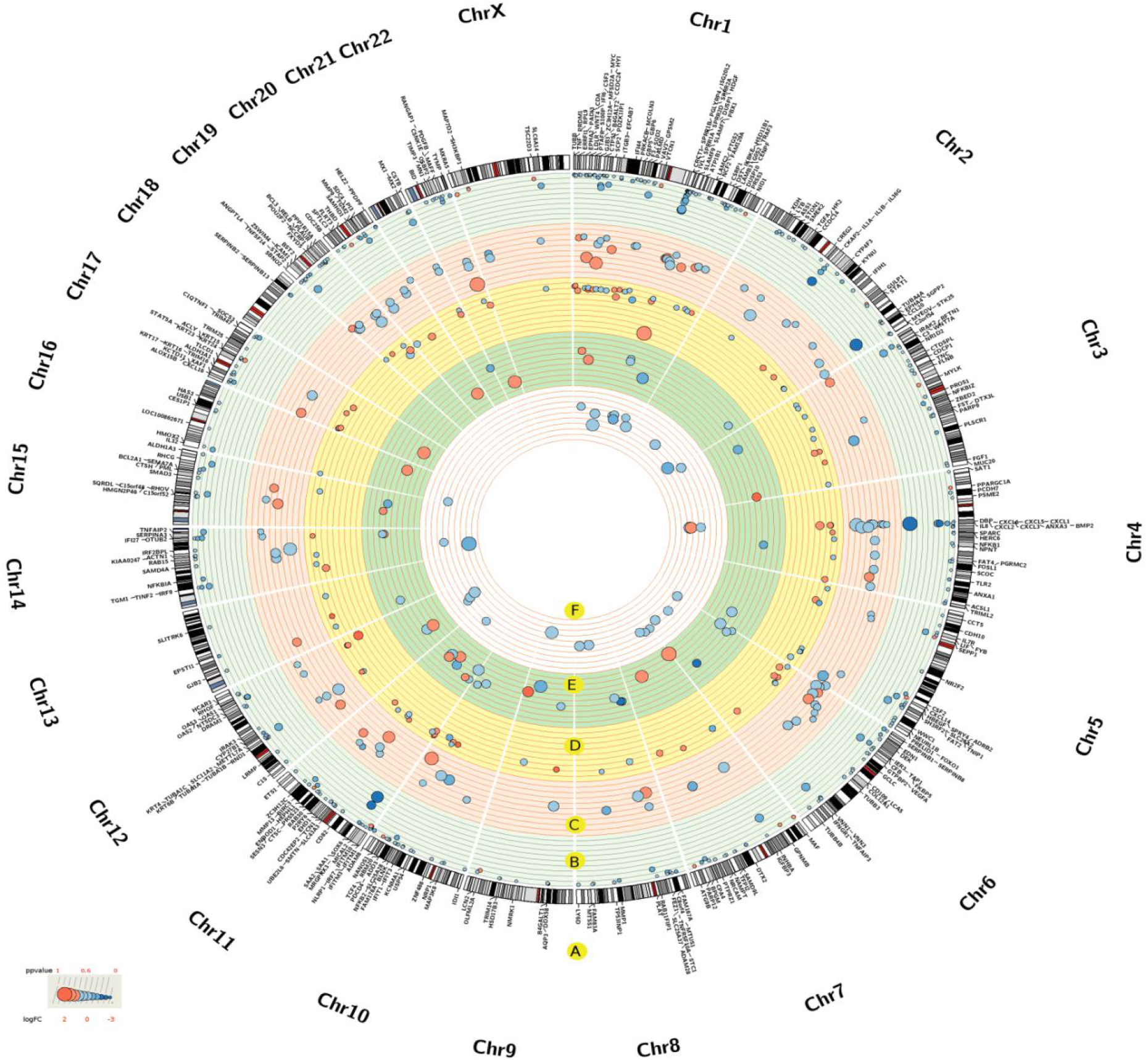
Significant DEGs across the five transcriptomic profiles, corresponding genes, chromosome locations, gene expression and significance scores. The DEGs related genes and chromosomal location (A). The DEGs information regarding host response to COVID-19 (B), Ebola (C), MERS-CoV (D), H1N1 (E) and SARS-CoV (F) viral infections. The pvalues were scaled were scaled across gene profiles according to maximum and minimum values (ppvalue). The circles size and color is linked to DEGs significance and gene expression (LogFC) scores, respectively.

**Figure 2 :**
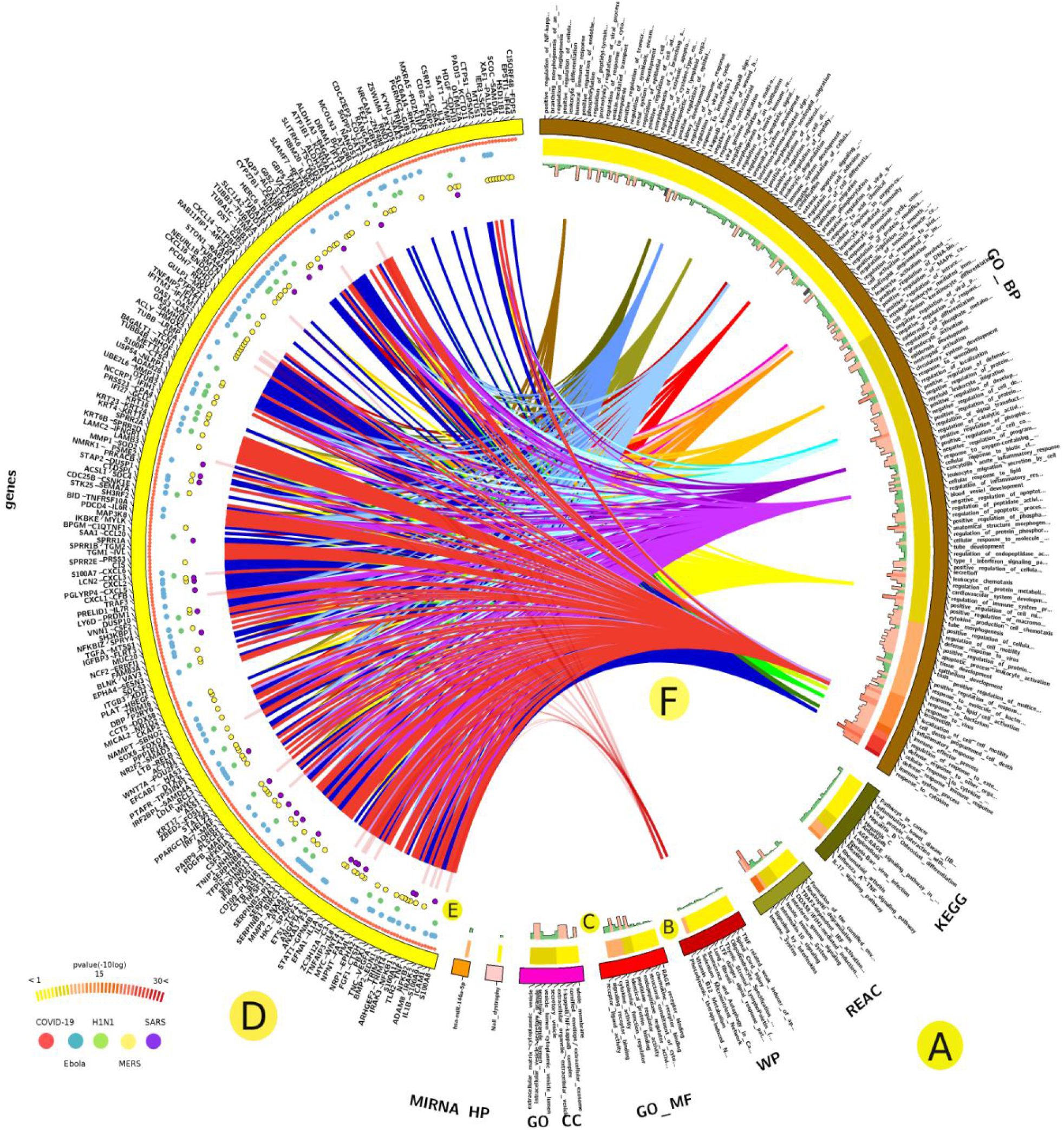
Analysis of the gene enrichment of DEGs correlated with the host response to COVID-19. Categories of GO terms (A), significance scores (−10log-pvalue) (B), and number of associated DEGs (C). The COVID-19-associated DEGs (D), status across the studied infectious diseases (E), and selected linked GO terms (F).

We particularly focused on the DEGs signature during COVID-19 infection and its overlap to other four viral infections. We found 173 DEGs are unique to COVID-19 (**Figure 3** **and Table S3**). Of these genes, *SAA2* was the most significant (−10logp-value of 81) (**Table S2**). GO analysis demonstrated that certain genes, such as *CSF3*, *CSF2, IL1B,* and *PTGS2*, are linked to the IL-17 signaling pathway, cytokine signaling in the Immune system, and have been reported in the host response to Rhinovirus infection (**Figure S1**). Overall, the terms of the biologic process, such as keratinocyte / epithelial cell differentiation, organ development, cell component movement and cell death, are very significant among these genes (**Table S4**), whereas molecular function such as RAGE receptor binding, cytokine activity, and metal ion binding are highly recognized (**Table S4**).

**Figure 3:**
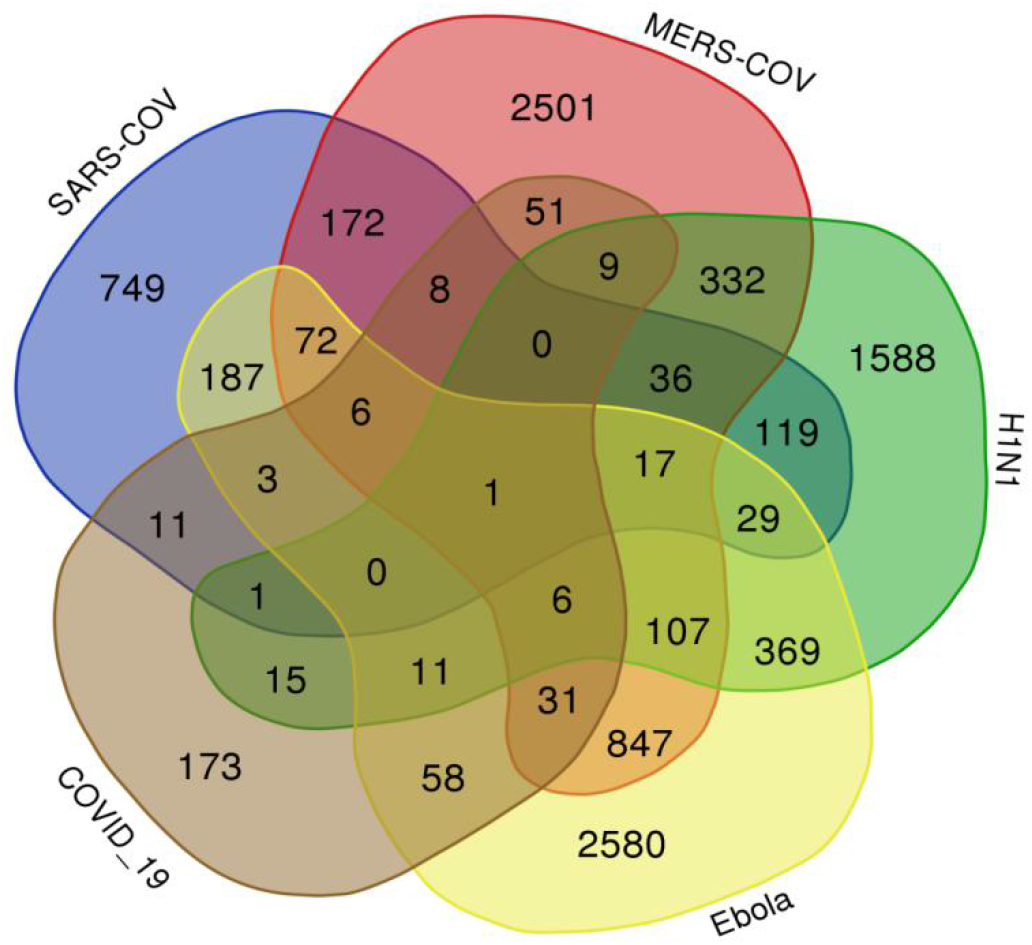
The Venn diagram of viral associated genes. The number of uniquely shared genes associated with the host response to COVID-19, Ebola, H1N1, MERS-CoV, and SARS-CoV viral infections.

Comparative gene expression analysis of the five viral infection (COVID-19, Ebola, H1N1, MERS-CoV, and SARS-CoV) yielded the *SERPINB1* as common response gene among the five infection. COVID-19 and Ebola uniquely shared 58 DEGs, followed by 51 DEGs between COVID-19 and MERS-CoV (**Figure 3 and Table S3**). Among the Ebola-shared *TNIP1, ICAM1,* and *CFB* genes were highly associated with COVID-19 (−10logpvalue > 40), while genes such as *TLR2, FOXO1*, and *MYC* were highly associated with cytokine response and cell death (**Figure S2** **and Table S2**). The GO molecular terms of these genes highlighted the biological functions of phospholipase inhibitor activity (including phospholipase A2), and heparin binding (including glycosaminoglycan). While biological processes such as cell surface receptor signaling pathways and cell death are highly significant (**Table S4**). The MERS-CoV-shared genes *KRT6B* and *TNFAIP3* have a high p-value associated with COVID-19, whereas genes such as *OAS1-3, IRF9, IRF7, STAT1, PML* and *IFIH1* are highly associated with host responses to viral infection and type I interferon (**Figure S3**). Biological processes related to virus response, Type I interferon signaling and the cytokine-mediated signaling pathway are highly redundant. While the biological functions of 2-5-oligoadenylate synthetase activity, double-stranded RNA binding, adenyltransferase activity, metal ion binding and related to growth activity, such as epidermal growth, are quite significant (**Table S4**). COVID-19, Ebola, and MERS-CoV share uniquely 31 genes, of which *BIRC3, MX1*, and *IL8* are strongly linked to COVID-19 (−logpvalue 23, 37, and 105, respectively) (**Figure 3**, **Table S2 and Table S3**). Among these genes, *DDX58* and *IFIT1* are highly associated with cytokine response, NF-kappa B signaling pathway, and immune response to virus infection (**Figure S4**).

The gene expression profile of COVID-19 signify genes such as *MX1, BIRC3, IRAK2, CXCL5, NRCAM, FGF1, MMP9, SAA1, LCN2, IFI27, TNFAIP3, OAS1, IL6, XAF1, IL8,* and *CXCL3* compared to Ebola, H1N1, MERS-CoV, and SARS-CoV. The host gene expression of these genes has changed exponentially relative to other infections studied (**Figure 1**, **S5** **and Table S5**). This list of genes are mostly related to IL-17 signaling pathway, TNF signaling pathway and host response against viral infection (**Figure S6**).

Analysis of gene enrichment showed that only three GO terms that are shared between COVID-19 and other viral infections (**Figure 4** **and Table S3**), including cellular component, protein binding and cytoplasm. COVID-19 was uniquely characterized by 535 GO terms, including stimulus response, cell communication, and defense response to bacterial infection (**Table S6**). COVID-19 shared 96 GO terms with Ebola, where GO terms related to the regulation of cell death are substantially shared. In addition, COVID-19 and MERS-CoV have uniquely shared 32 GO terms, most of which are linked to cell defense against viral infection and immunity, and metal ion response (**Figure 4**, **Table S3 and S6)**.

**Figure 4:**
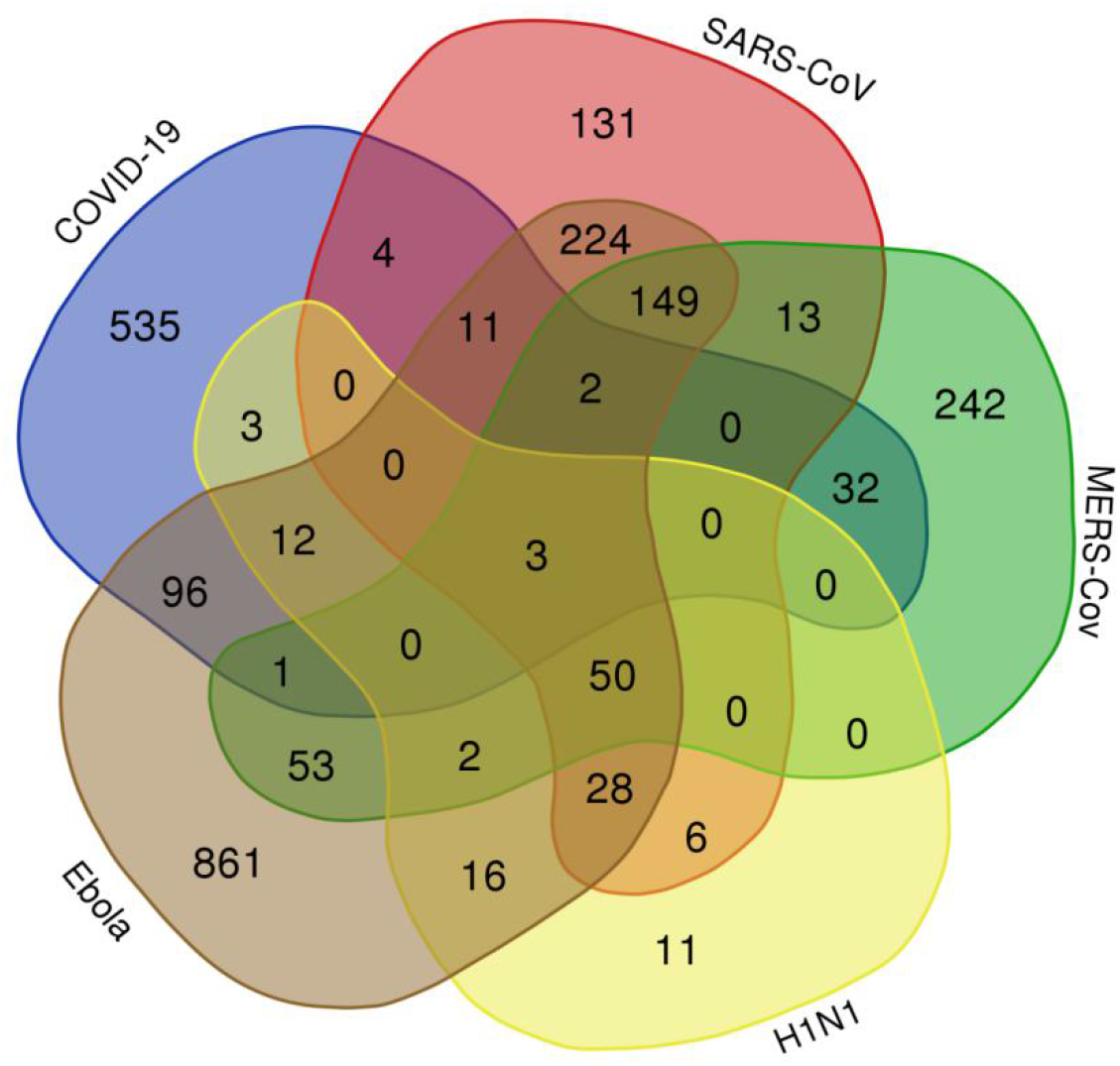
The Venn diagram of viral associated GO terms. The number of uniquely shared GO terms of DEGs associated with the host response across COVID-19, Ebola, H1N1, MERS-CoV, and SARS-CoV viral infections.

We used the PPI association network analysis to identify the shared DEGs between COVID-19 and the other four viral infections (**Figure 5**). The PPI network signify genes such as *IL6, TNF, IL8, VEGFA, IL1B, MMP9, STAT1, TLR1, CXCL1, ICAM1, TLR2*, and *IRF7* with high interaction activity. Some genes are associated with both COVID-19 and Ebola, and a few are shared with MERS-CoV (**Figure 5**). The PPI analysis and gene enrichment analysis of these hyper-interactive genes showed significant biological functions connected to rheumatoid arthritis, AGE-RAGE signaling pathway, malaria, hepatitis B, and influenza A (**Figure 6**).

**Figure 5:**
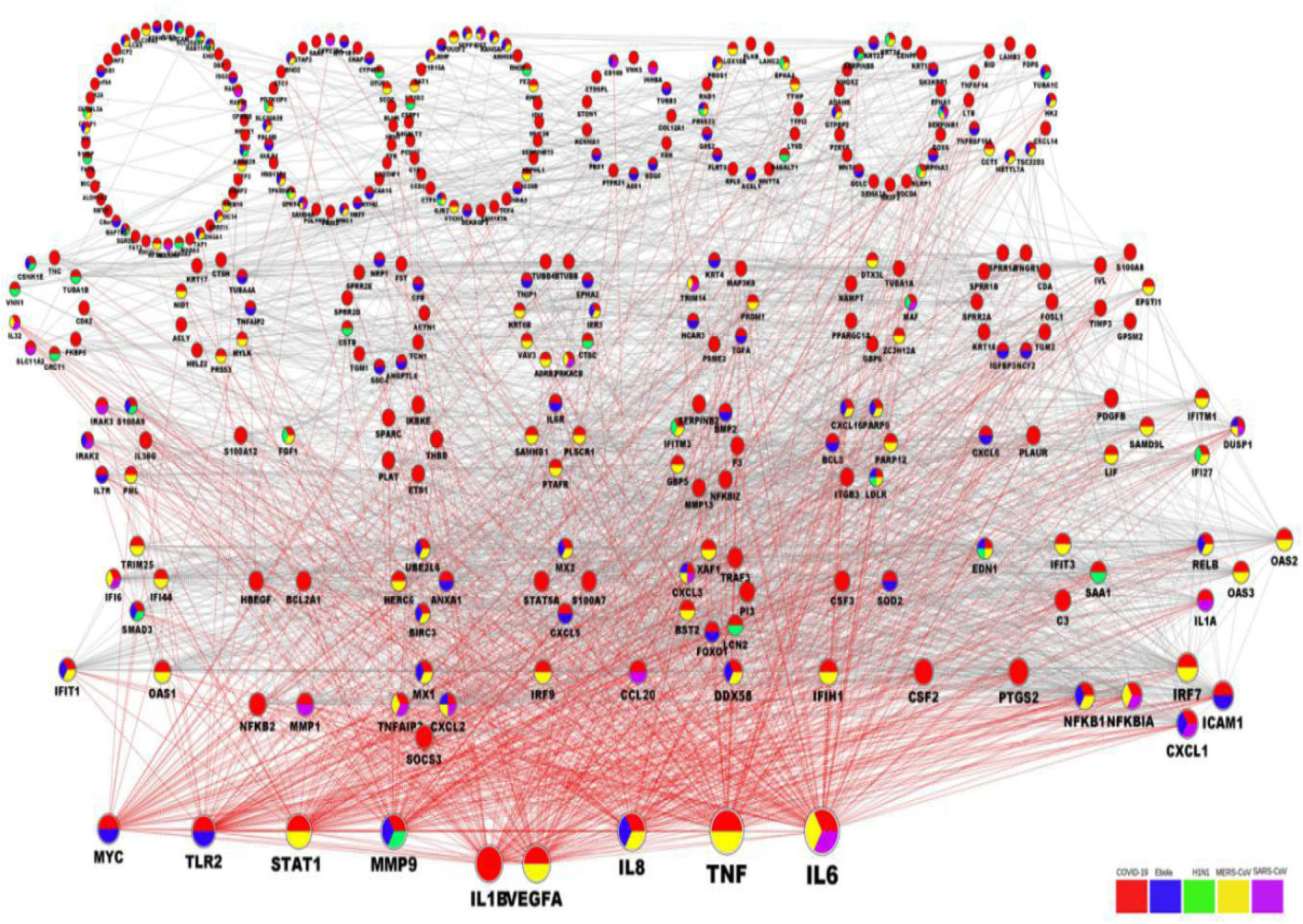
The PPIs network of DEGs associated with COVID-19. The PPI of host expressed DEGs under COVID-19 infection. DEGs shared between COVID-19 and Ebola, H1N1, MERS-CoV, and SARS-CoV are color-coded according to kind of infection. The gene node size is relative to its interaction activity. DEGs are collected in different groups according to their level of interaction activity.

**Figure 6:**
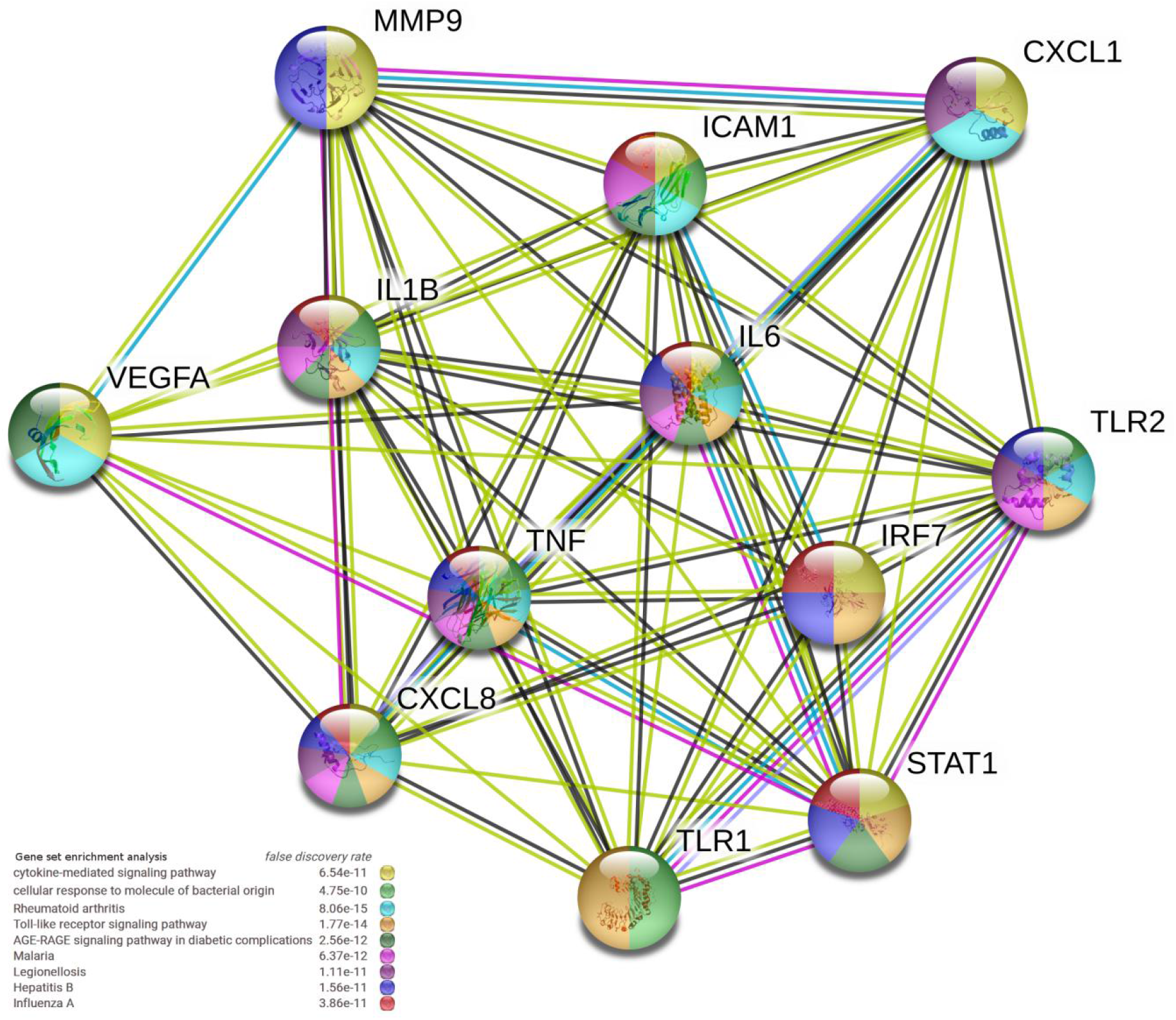
The PPIs network and gene enrichment analysis of highly interactive genes associated with COVID-19.

## Discussion

This study mainly aimed to determine the unique host gene expression signature response to COVID-19 infection compared to SARS-CoV, MERS-CoV, Ebola, and H1N1, which will help us to understand the differences and similarities in host responses to various respiratory viruses. To our knowledge, this is the first study to perform such a transcriptomic comparison between these five viral infections. We focused on mapping the potential biological pathways and GO enrichment that are more specific to COVID-19.

The analysis of the host DEGs through COVID-19 infection highlighted the role of *SAA2, CCL20,* and *IL8* genes (**Figure 1B** **and Table S2**). Recently, this link between the serum amyloid A 2 (*SAA2*) gene and the COVID-19 infection has been proposed as a biomarker to differentiate the severity and prognosis of the COVID-19 infection. *SAA2* is an inflammation factor that has demonstrated its effectiveness as a sensitive indicator of clinical diagnosis (Li et al. 2020). In addition, we have observed the uniqueness of the *SAA2* gene expression in the COVID-19 infection relative to SARS-CoV, MERS-CoV, Ebola, and H1N1 viral infections (**Figure 3** **and Table S3**), which indicates its role in host response. On the other hand, *CCL20* gene has been related to lung carcinoma, where it controls proliferation and cell migration via the PI3 K pathway (Wang et al. 2016). These mechanisms are among the most important in host defense. Multiple genes belong to the interleukin gene family were identified in this study, such as *IL6, CXCL1, 3* and *5*, and the *IL-17* which have a significant association with the host response of COVID-19 (**Figure 1** **and Table S2**). In addition, *IL8* gene, which has been related to immune stimulus and a recognized locus of susceptibility to a specific respiratory virus (Hull et al. 2001). Such genes serve as key factors for controlling the growth of endothelial cells (Martin et al., 2009).

GO-based gene enrichment analysis demonstrated that many biological processes are closely related to the immune response (**Figure 2A**), including myeloid cell activation and neutrophil activation (**Figure 4C and 4E**). Interestingly, miRNAs-related gene pathway was overexpressed as a response to COVID-19 infection, which is known to play an important role against viral infection (Nur et al. 2015). Activation of miRNAs as a defense mechanism during lung infection could be related to its important role in physiological and pathological processes in the lung (Tomankova et al., 2010). Studying such a process could open a new way for treatment of COVID-19.

We identified a strong association between COVID-19 infection and GO related to Nuclear Factor Kappa-B (NF-kB) signaling and Tumor Necrosis Factor (TNF) signaling pathways (**Figure 2B**). The NF-kB pathway is closely related to pro-inflammatory and pro-oxidant responses, and is involved in the inflammatory responses in acute lung injuries. The regulation of NF-kB activation was proposed as a potential adjuvant treatment for COVID-19 infection (Zhang et al. 2020). TNF receptors are mainly involved in the inflammation and apoptosis; interestingly the interactions between viral proteins and intracellular components downstream of the TNF receptors demonstrated viral mechanism to evade the immune response (Herbein and O’brien 2000).

Among the genes that are unique in the host response to COVID-19 are *CSF2/3, and PTGS2*, which known to be involved in the immune responses against Rhinovirus infection (**Figure S1**). The relation between prostaglandin-endoperoxide synthase 2 (*PTGS2*/*COX-2*) gene and host response to COVID-19 infection could be due its role to down-regulate NF-κB mediated transcription, which is a critical element in some virus replication such as HIV-1 (Feistritzer and Wiedermann 2007). It was proposed that this gene is incorporated in the host immune response system against viral infection (Whitney et al., 2011). The colony-stimulating granulocyte factor (*G-CSF)* can alter the function of T-cells and induces Th2 immune response (Franzke et al. 2003). There is also some evidence of a link between elevated G-CSF expression level and the induction of the cellular immune response in H1N1 infected individuals (Sadeghi et al. 2020).

The GO-associated molecular function in COVID-19 host response yielded terms such as receptor for advanced glycation endproducts (*RAGE*) and metal ion binding (**Figure 2B** **and Table S4**). RAGE is highly expressed only in the lung, and is rapidly growing at inflammatory sites, primarily in inflammatory and epithelial cells. The triggering and upregulation of RAGE by its ligands correlate with increased survival rates (Sparvero et al. 2009). Additionally, RAGE has a secretory isoform that can have an independent causative effect on community-acquired pneumonia, such as pandemic influenza (H1N1) (Narvaez-Rivera et al. 2012). Although there is no evidence to link this to COVID-19 infection, it is worth further investigation.

The host response to the five viruses shared the plasminogen activator (*SERPINB1*) as a common gene signature (**Figure 3** **and Table S3**). This gene is highly correlated with lung chronic airway inflammation such as asthma (Dijkstra et al. 2011). The *SERPINB*1 acts in host-pathogenic interactions and possesses some antiviral activity across infections of rhabdovirus, hepatitis C, and influenza A (Dittmann et al. 2015) (Estepa and Coll 2015).

Among the five viral infections, we found that GO terms were mostly enriched between COVID-19 and Ebola (**Figure 4** **and Table S3**). Such overlap suggested certain genes and gene families, which could explain the aggressiveness of COVID-19 infections. Within these GO enriched pathways, the *TNIP1, ICAM1,* and *CFB* were most significantly associated with COVID-19 (logpvalue > 40) (**Figure 1** **and Table S2)**. The *TNIP1* gene encodes the A20-binding protein that plays a role in autoimmunity and tissue homeostasis by controlling the activation of the kappa-B nuclear factor (Bowes et al. 2011). *TNIP1* reduction sensitizes keratinocytes to post-receptor signaling after interaction to *TLR* agonists and has the ability to activate immune cells and induce inflammation (Kaczanowska et al. 2013). The correlation between *TNP1* and COVID-19 (**Figure 1** **and Table S2**) could be due to its role in suppressing NF-kB pathway and therefore regulating the overexpression of viral proteins (Nimmerjahn et al. 2004) (Ramirez, Gurevich, and Aneskievich 2012). The *ICAM-1* intercellular adhesion molecule plays a major role in the infectivity and neutralization of the HIV-I and controls the survival of the influenza virus in lung epithelial cells during the early stages of infection (Othumpangat et al. 2016). Forkhead Box O1 (*FOXO1*) is a transcription factor that plays an important role in the regulation of insulin signaling for gluconeogenesis and glycogenolysis. There has been a strong relationship between *FOXO1* and viral infections. The *FOXO1* binds hepatitis B virus DNA and activates its transcription (Shlomai and Shaul 2009). The *FOXO1* reported to negatively regulate cellular antiviral response by promoting degradation of interferon regulatory transcription factor 3 (*IRF3*) (Lei et al. 2013). In addition, it was reported that differentiation of CD8 memory T cells depends on *FOXO1*, where it plays an intrinsic role in the establishment of a post-effector memory program which is important for the formation of long-lived memory cells capable of immune reactivation (Michelini et al. 2013).

GO analysis of genes uniquely shared between COVID-19 and Ebola highlighted the activity of the inhibitor of phospholipase, in particular phospholipase A2 (PLA2) (**Figure 4** **and Table S4).** Interestingly, synthetic and natural PLA2 inhibitors have been a viable treatment of oxidative stress and neuroinflammation connected with neuropathogenic disorders (Ong et al. 2015). Such lipid mediators are considered to play a major role in diseases associated with cancer and inflammation such as arthritis, allergy and asthma (Greene et al. 2011). Some reports suggested a potential link between PLA2-generated lipid mediators and viral infection, where these infection alters the lipid mediators of this pathway to initiate infection and pathogenesis (Chandrasekharan and Sharma-Walia 2019). Given the important association between heparin-binding GO and activation of T cells against virus infections like influenza (Skidmore et al. 2015), their interaction with COVID-19 infection has not been documented. In comparison, glycosaminoglycan-binding molecules are essential for the action of certain *in vivo* chemokines. Some glycosaminoglycans are required for respiratory syncytial viral infection and are important for the entry of a bacterial pathogen into the biological system (Chang et al. 2011). Some oncofetal antigens which target such proteins were used to control parasites of malaria (Mette et al., 2019). This might support any of the recent suggestion of using pharmaceuticals derived from glycosaminoglycan to control the infection with COVID-19 (Favaloro and Lippi 2020).

MERS-CoV uniquely shared 51 DEGs with COVID-19 (**Figure 3** **and Table S3**). Among the most significant shared genes that are associated with COVID-19 are *KRT6B and TNFAIP3*. Keratin 6B (*KRT6B*) is a type II cytokeratin, which is an important biomarker for lung adenocarcinoma (Xiao et al. 2017). These genes are known as a virus-induced host factors that control the recruitment of T-cells and correlates to chronic virus infections (Wang et al. 2020). In addition, the tumor necrosis factor, alpha-induced protein 3 (*TNFAIP3*), is a central regulator of immunopathology and associated with the maintenance of immune homeostasis and severe viral infections (Mérour et al. 2019; Li et al. 2017).

We identified many DEGs that are classified as “antiviral genes” that are shared between MERS-CoV and COVID-19 (**Figure 3**). Most of these DEGs are associated with host response to virus infection, and type I interferons (**Figure S3**). This high number of DEGs could indicate their potential role in host defense against COVID-19 infection. For instance, the regulation of *OAS1-3* is highly correlated with host response to viral infections (Melchjorsen et al. 2009). While genes such as *IRF9*, *PML, IRF7, STAT1* and *IFIH1* are related to interferon signaling (Ramana et al. 2002).

COVID-19, Ebola, and MERS-CoV shared uniquely 31 genes, of which, *BIRC3* and *MX1* are highly linked to COVID-19 (**Figure 3** **and Table S3**). The Baculoviral IAP Repeat Containing 3 (*BIRC3*) is associated Marginal Zone B-Cell Lymphoma, Lymphoma, and was suggested as a novel NK cell immune checkpoint in cancer (Ivagnès et al. 2018). While MX Dynamin Like GTPase 1 (*MX1*) is an interferon-inducible protein that associated with viral infections of Influenza and Viral Encephalitis (Ciancanelli et al. 2016). The link between the gene expression of *BIRC3* and *MX1* have been hypothesized as a part of small group of genes controlling host response against viral infections, including Human Herpes Virus type 6Α (HHV-6Α) infection (Rouka 2018). Additionally, *Mx1* protein contributes to the novel antiviral activity against classical swine fever virus (Chen et al. 2020). Among genes that are uniquely shared between COVID-19, Ebola, and MERS-CoV, interferon Induced Protein With Tetratricopeptide Repeats 1 (*IFIT1*) and DExD/H-Box Helicase 58 (DDX58) which high a significant potentiality (**Figure S4**). Recently, the uniqueness of *DDX58* gene expression under COVID-19 viral infection has been reported (Blanco-Melo et al. 2020). *IFIT1* plays a crucial role in some viral infections, where Hepatitis E virus polymerase binds to *IFIT1* to shield the viral RNA from translation inhibition mediated by *IFIT1* and enhances the interferon response in murine macrophage-like cells (Mears et al. 2019) (Pingale, Kanade, and Karpe 2019).

The COVID-19 gene expression profile demonstrated multiple genes in conjunction with Ebola, H1N1, MERS-CoV, and SARS-CoV (**Figure 3**, **S6** **Table S5**). Most of these genes are linked to the viral infection immune response of the host, except for genes such as *FGF1* and *NRCAM*. The Neuronal Cell Adhesion Molecule (*NRCAM*) is related to neurological diseases such as Alzheimer (Brummer et al. 2019). Significant NRCAM gene expression has been observed under specific circumstances, such as neuroinflammation triggered by influenza A long-term viral infection (Hosseini et al. 2018). *FGF1*, also known as acidic fibroblast growth factor (aFGF), is a cellular growth factor and signaling protein encoded by the FGF1 gene. *FGF1* is a strong angiogenic factor controls the development of new blood vessels (Marwa et al. 2016) and has been detected through studying endothelial cells infected with influenza virus (Zeng et al. 2012).

The PPI analysis highlighted the genes COVID-19 shared with other viral infections that have high interaction activity (**Figure 5**). By selecting high interactive genes, we used an analysis of gene enrichment and PPI to identify more information about the function of these genes. It was clear from the results that the genetic pathways associated with Rheumatoid arthritis, AGE-RAGE signaling pathway, Malaria, Hepatitis B, and Influenza A were of high significance (**Figure 6**). The correlation between host response to Rheumatoid arthritis, Malaria and COVID-19 has been mysterious to date. Despite the fact that several Rheumatoid arthritis and malaria drugs are available, with some efficacy against COVID-19 infection (Favalli et al. 2020) (Gao, Tian, and Yang 2020). Our results suggest that the link between these diseases and the infection with COVID-19 is more related to PPI interactions. In addition, the PPI network has shown that these genes are highly significant across other infectious diseases such as Ebola, MERS-CoV and SARS-CoV.

## Conclusion

We compared five transcriptomic profiles for cell host infection with COVID-19, Ebola, H1N1, MERS-CoV and SARS-CoV. Our analysis identified several key aspects of host response to COVID-19 infection where essential immunity genes and biological pathways could be used for understanding the pathogenesis of COVID-19 infection. Common and specific genetic factors and pathways have been identified that characterize the immune pathology of COVID-19 infection. Our research outlined the relationship between Ebola's cellular host response and COVID-19, where many genes and GO words are enriched. Genes related to immune regulation, including FGF1 and *FOXO1*, and those associated with extreme inflammation, such as *NRCAM* and *SAA2*, have been closely associated with cellular response to COVID-19 infection. In addition, common interleukin family members, in particular IL-8, IL-6, demonstrated a special relationship with COVID-19 infection, indicating their key importance. The GO evaluation highlighted pathways for RAGE, miRNA and PLA2 inhibitors, which were first identified in this study as possible pathways highly associated with the host response to COVID-19 infection. Some of these pathways, such as PLA2 inhibitors, may hold the key for potential drugs to manage COVID-19 infections. The PPI study sheds light on genes with high interaction activity that COVID-19 shares with other viral infections, where the findings showed that the genetic pathways associated with Rheumatoid arthritis, AGE-RAGE signaling system, Malaria, Hepatitis B, and Influenza A were of high significance. Our work also shows that the combination of different types of experimental methods and parameters have been effective in studying the etiology of COVID-19 immunopathology compared to similar viral infections. In this regard, further research in this direction will be promising for characterizing new diagnostic biomarkers in the future or as surrogates for assessing the effectiveness of potential innovative therapies.

## Data availability

All data are freely available at https://doi.org/10.5281/zenodo.3783510

## Supplemented Tables

**Table S1 :** The data information used in this study.

**Table S2**: The information of DEGs associated the host response of COVID-19, Ebola, H1N1, MERS-CoV, and SARS-CoV viral infections.

**Table S3**: The Venn analysis results of DEGs and GO terms uniquely shared across of COVID-19, Ebola, H1N1, MERS-CoV, and SARS-CoV viral infections.

**Table S4**: Selected gene enrichment analysis of uniquely shared group of genes across the host response of COVID-19, Ebola, H1N1, MERS-CoV, and SARS-CoV viral infections.

**Table S5**: The gene expression information of DEGs that COVID-19 share with the studied infectious diseases.

**Table S6**: Selected gene enrichment analysis of uniquely shared group of GO terms across the host response of COVID-19 and studied viral infections.

**Figure S1 :**
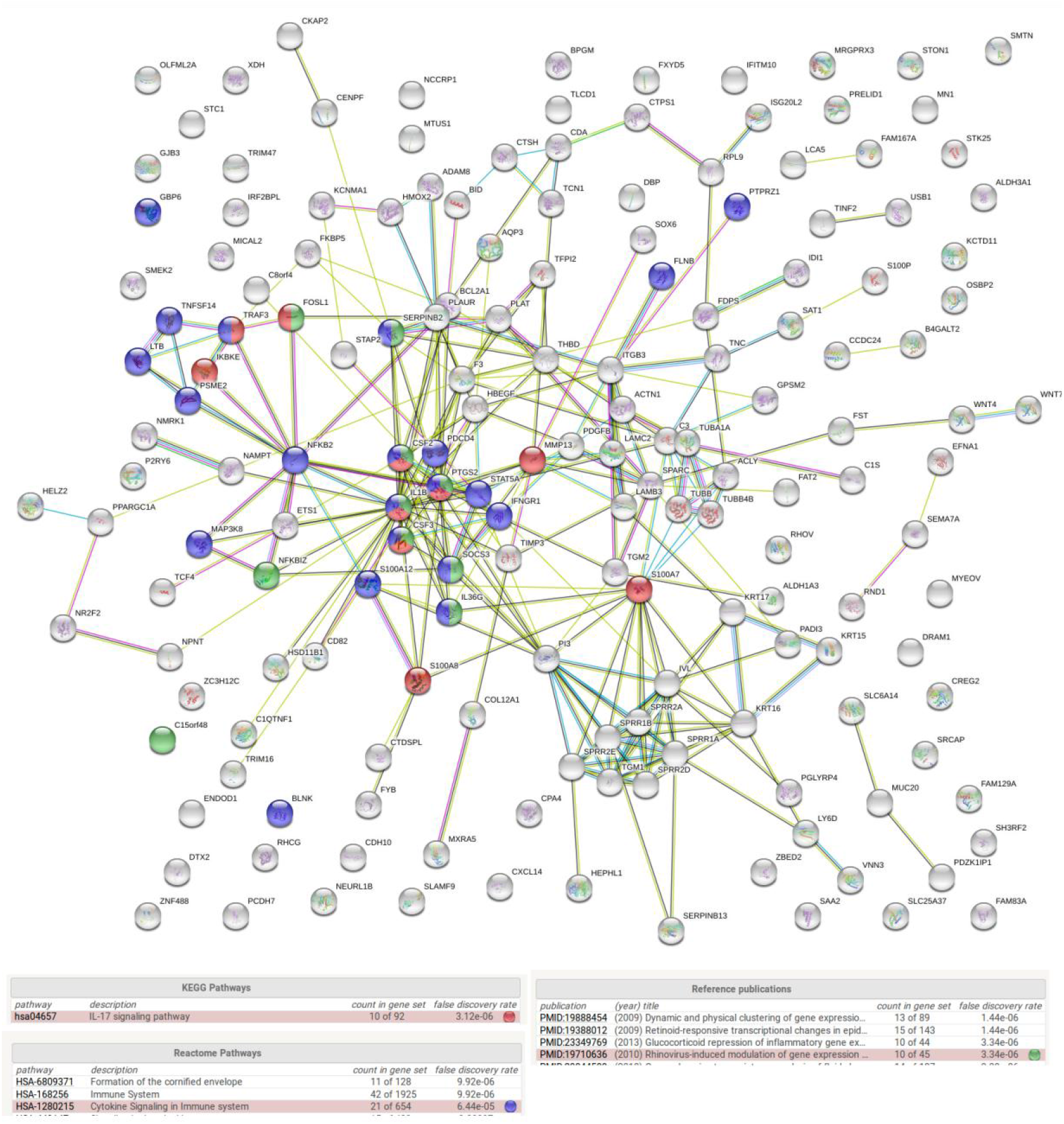
The PPI network and gene enrichment analysis of the 173 genes that characterized the host response of COVID-19.

**Figure S2:**
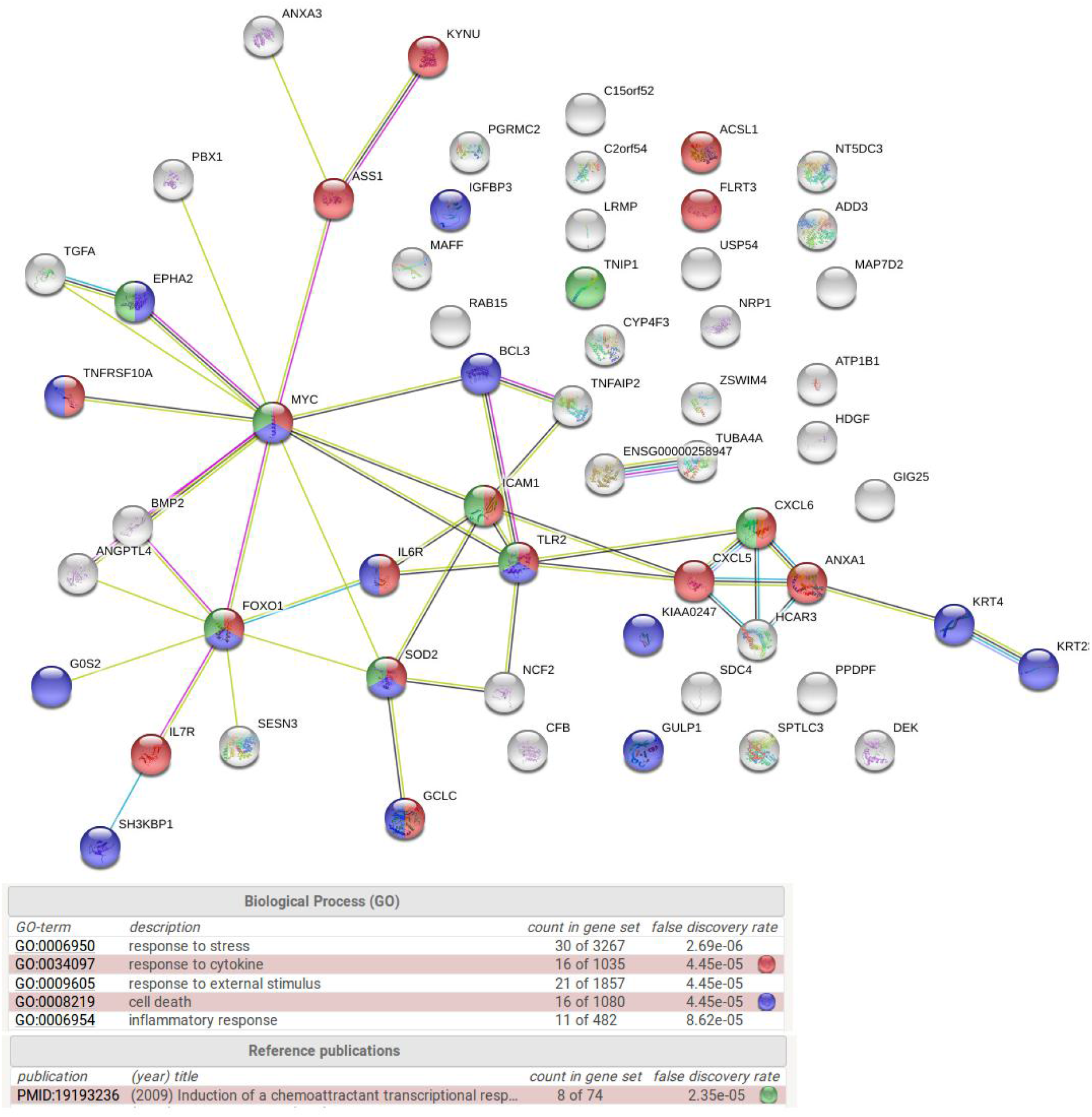
The PPI network and gene enrichment analysis of the 58 genes that are uniquely shared between COVID -19 and Ebola viral infections.

**Figure S3 :**
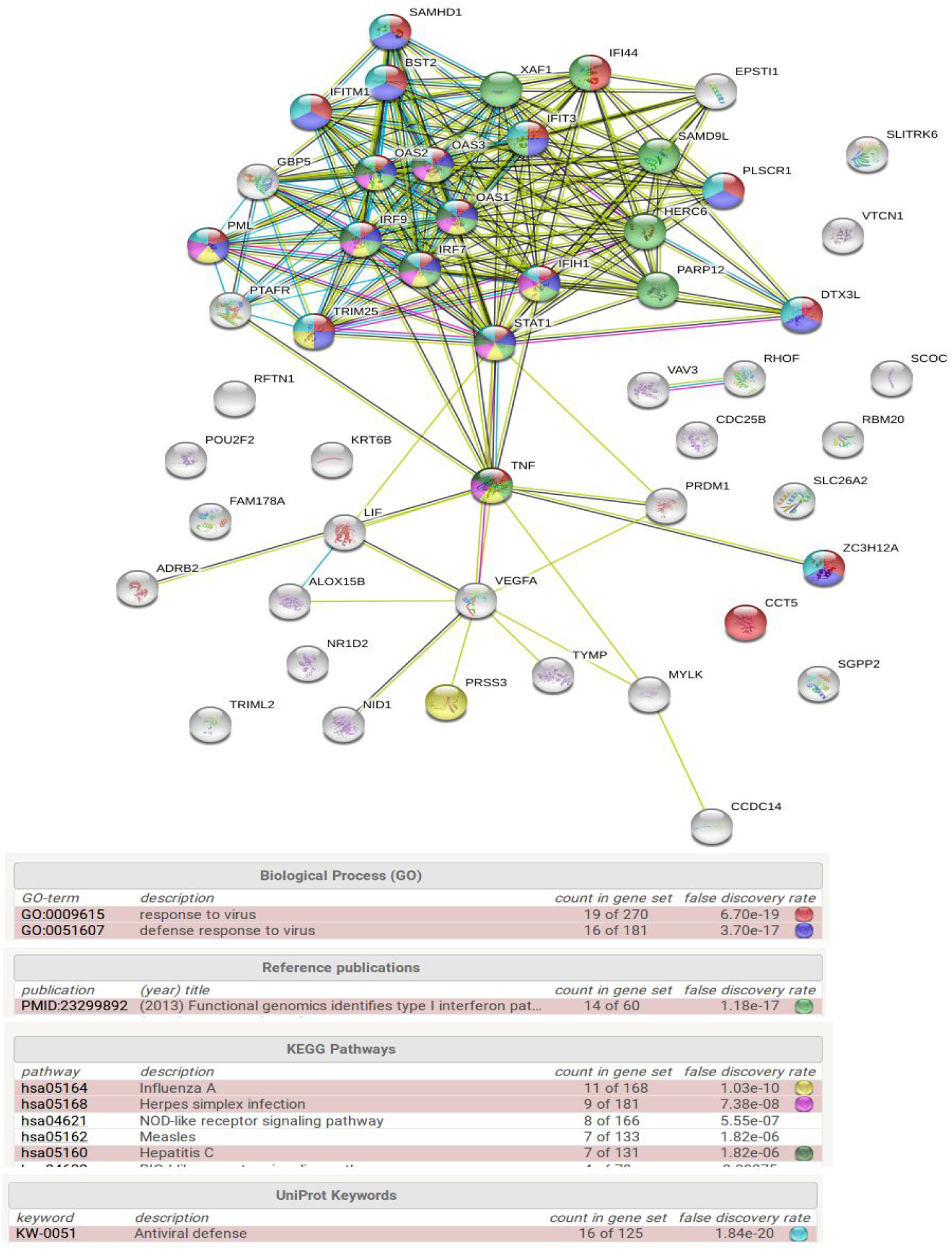
The PPI network and gene enrichment analysis of the 51 genes that are uniquely shared between COVID-19 and MERS-CoV viral infections.

**Figure S4 :**
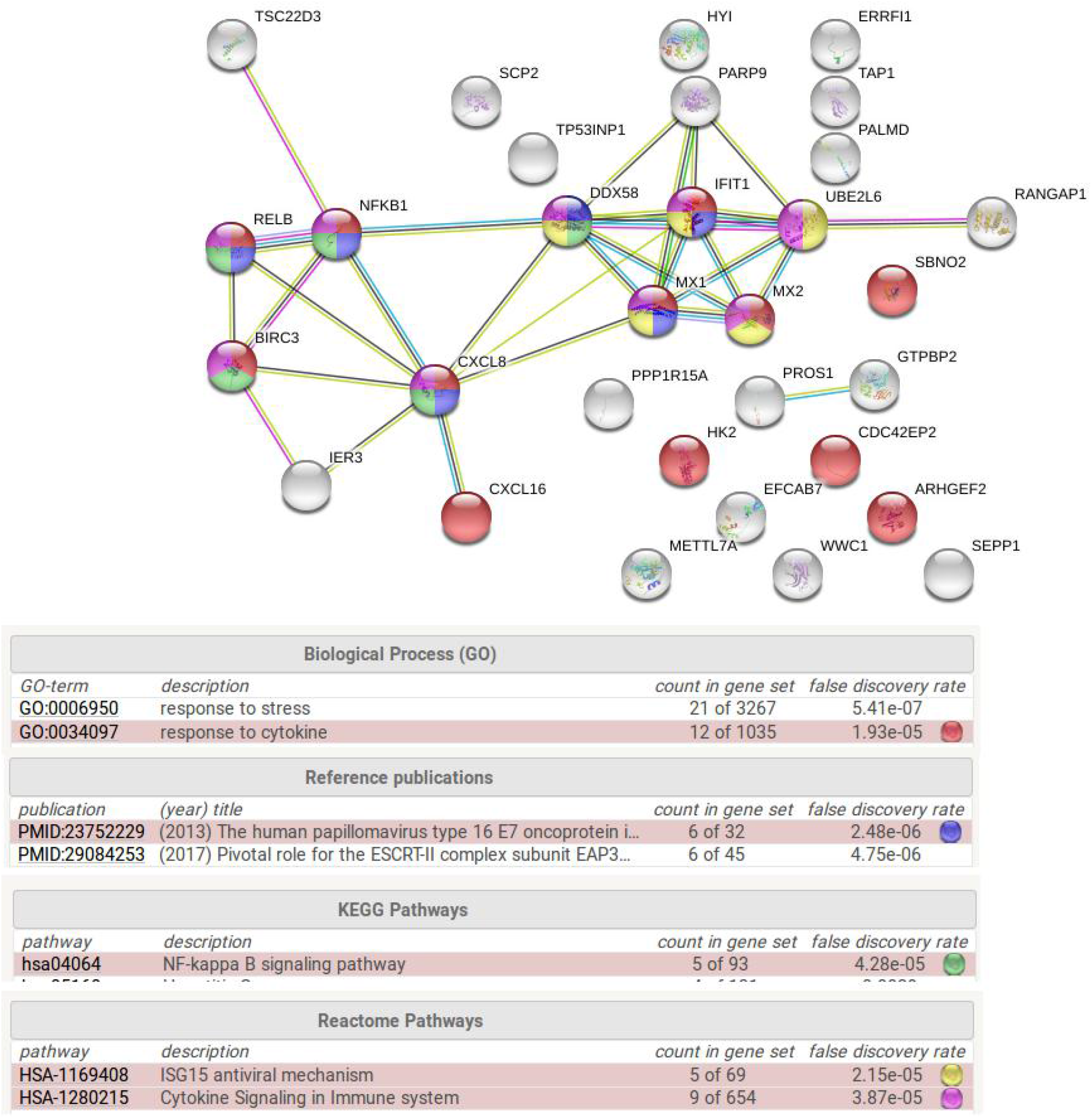
The PPI network and gene enrichment analysis of the 31 genes that are uniquely shared between COVID-19, Ebola, and MERS-CoV viral infections.

**Figure S5:**
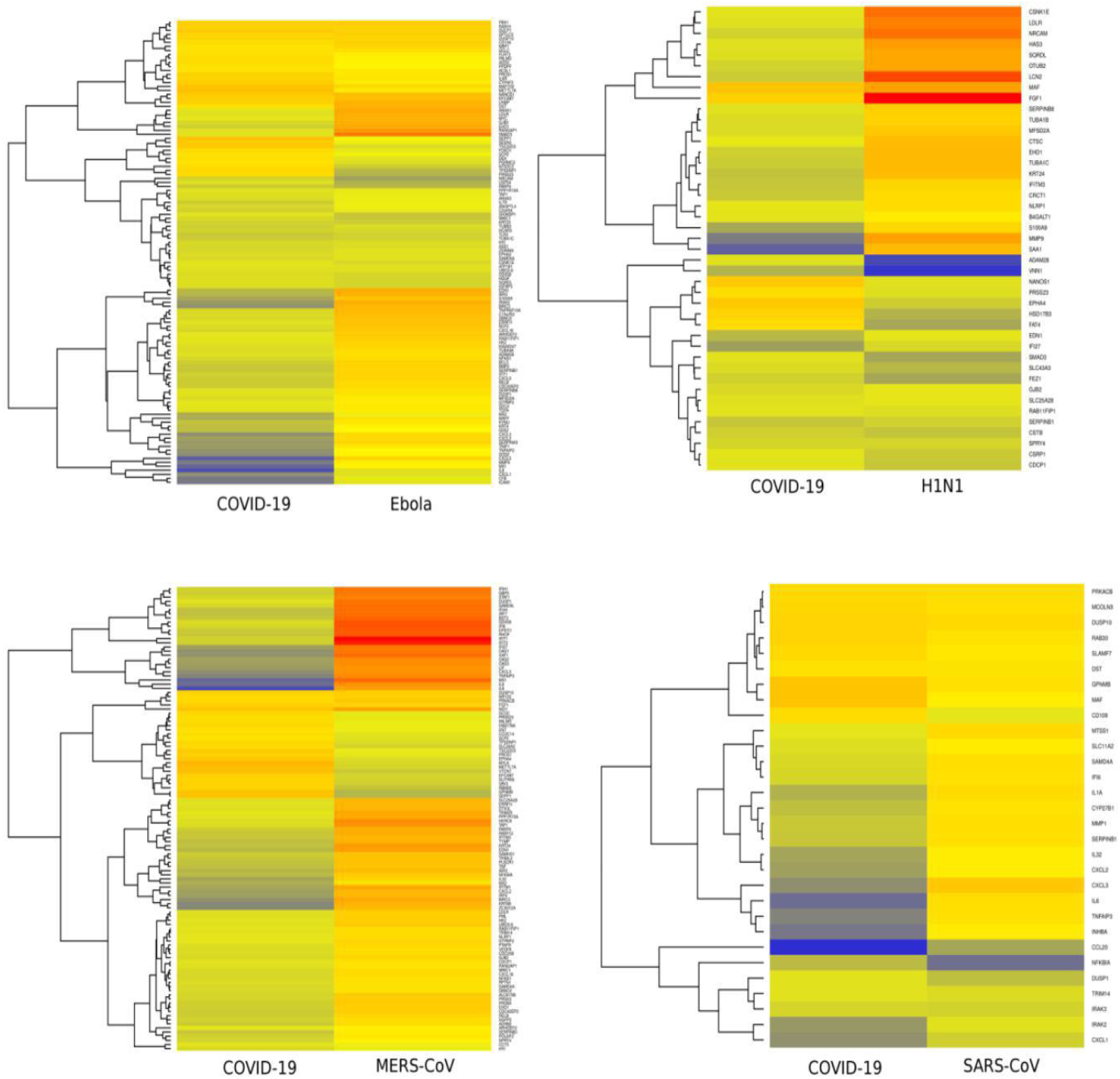
The gene expression heatmap of genes COVID-19 shares with different viral infections.

**Figure S6 :**
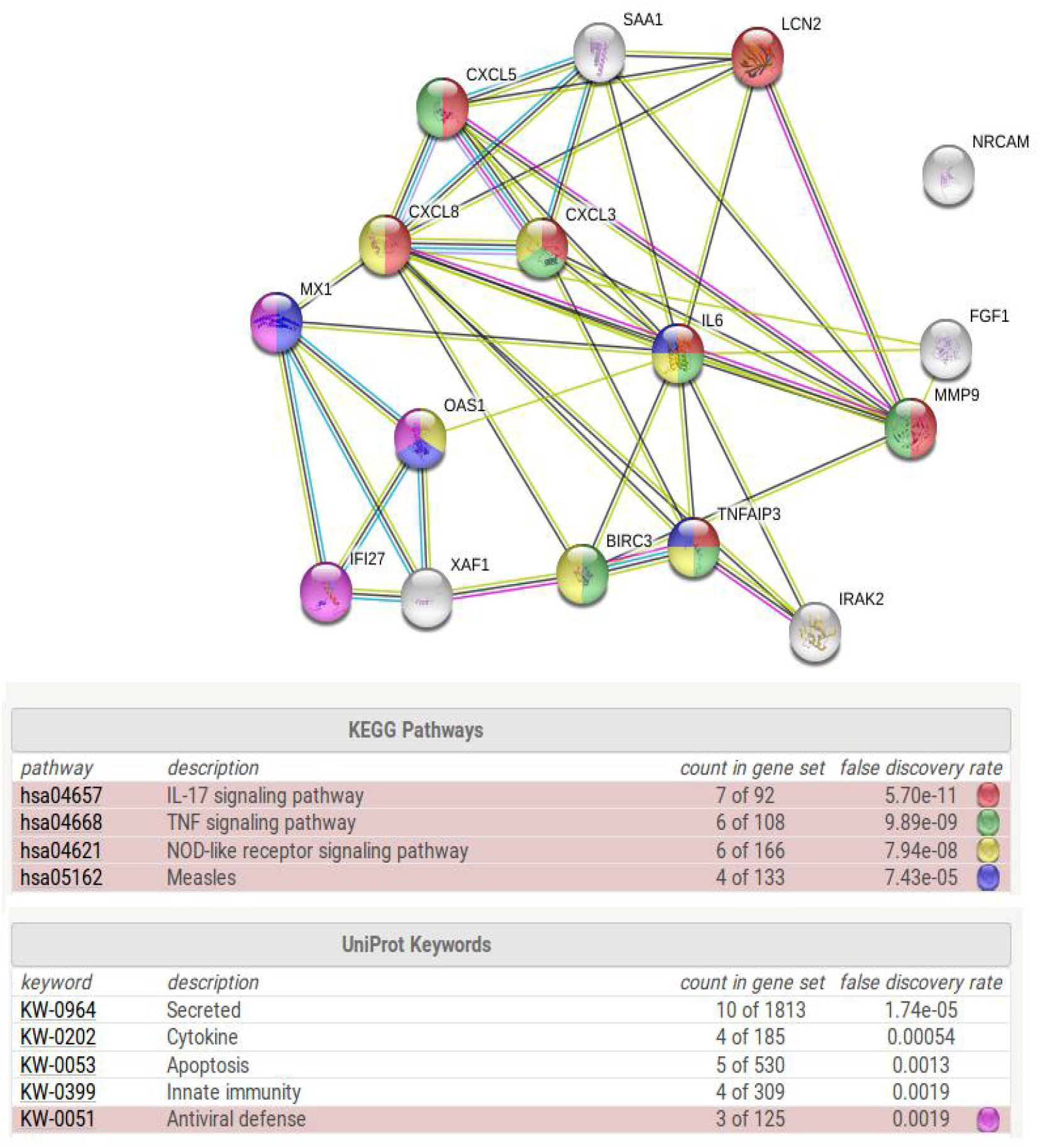
The PPI network and gene enrichment analysis of genes that are differentially expressed across studied viral infections and shared with COVID-19.

